# Mechanisms of Neural Representation and Segregation of Multiple Spatially Separated Visual Stimuli

**DOI:** 10.1101/2025.09.11.675659

**Authors:** Steven Wiesner, Bikalpa Ghimire, Xin Huang

## Abstract

Segregating objects from one another and the background is essential for scene understanding, object recognition, and visually guided action. In natural scenes, it is common to encounter spatially separated stimuli, such as distinct figure-ground regions, adjacent objects, and partial occlusions. Neurons in mid- and high-level visual cortex have large receptive fields (RFs) that often encompass multiple, spatially separated stimuli. It is unclear how neurons represent and segregate multiple stimuli within their RFs, and the role of spatial cues for such representation. To investigate these questions, we recorded neuronal responses in the middle temporal (MT) cortex of monkeys to spatially separated stimuli that moved simultaneously in two directions. We found that, across motion directions, response tuning to multiple stimuli was systematically biased toward the stimulus located at the more-preferred RF subregion of the neuron. The sign and magnitude of this spatial-location bias were correlated with the spatial preference of the neuron for single stimuli presented in isolation. We demonstrated that neuronal responses to multiple stimuli can be captured by an extended normalization model, which is a sum of the responses elicited by individual stimuli weighted by the spatial preference of the neuron. We also proposed a circuit implementation for the model. Our results indicate that visual neurons leverage spatial selectivity within their RFs to represent multiple spatially separated stimuli. The spatial-location bias in neuronal responses enables individual components of multiple stimuli to be represented by a population of neurons with different spatial preferences, providing a neural substrate for segregating multiple stimuli.

**Significance Statement:** Elucidating how neurons represent multiple visual stimuli is crucial for understanding the principles and mechanisms of neural coding. We found that the neuronal response in MT to spatially separated moving stimuli can be captured by the well-known normalization model, with an important new extension: the responses elicited by the individual stimulus components are combined and weighted by the neuron’s spatial preference for single stimuli within its receptive field. Consequently, the response of a neuron to multiple stimuli can be substantially biased toward the stimulus at the neuron’s preferred spatial location. Our results revealed a previously unknown coding strategy for representing and segregating multiple spatially separated stimuli. Our proposed circuit implementation provides insight into the neural mechanisms underlying spatial preference-weighted normalization.

## Introduction

In natural vision, it is rare to encounter an isolated object presented on a blank background. Instead, natural scenes are often complex and contain multiple entities. The ability to segregate objects from one another and from their background is central to scene interpretation, object recognition, and visually guided action.

Visual motion offers potent cues for scene segmentation that the visual system readily exploits (Wertheimer, 1938; Braddick, 1993; Stoner & Albright, 1993; Britten, 1999; Wagemans et al., 2012). The extrastriate area MT in primates serves as a key hub for processing motion information and plays a crucial role in motion-based segmentation (Born & Bradley, 2005; Britten, 2003; Pasternak & Tadin, 2020). Previous neurophysiological studies have investigated how MT neurons represent and segregate overlapping moving stimuli in their receptive fields (RFs) that give rise to the perception of transparent motion (Snowden et al., 1991; Stoner & Albright, 1992; Qian & Andersen, 1994; Treue et al., 2000; Xiao et al., 2014; McDonald et al., 2014; Xiao & Huang, 2015; Wiesner et al., 2020; Chakrala et al., 2024; Huang et al., 2025). Our laboratory has found that the responses of MT neurons to overlapping moving stimuli often preferentially represent one stimulus component with either a higher luminance contrast or motion coherence (Xiao et al., 2014), one of two competing motion directions (Xiao & Huang, 2015), speeds (X. Huang et al., 2025), or binocular disparities (Chakrala et al., 2024). These studies suggest a common pattern: neurons tend to employ a strategy of biased mixing of stimulus components, rather than equally pooling them. This strategy enables neurons to better represent individual stimulus components.

Using transparent moving stimuli has the advantage of isolating motion cues from spatial cues, as stimulus components are superimposed and occupy the same spatial region. However, in natural environments, it is more common to encounter spatially separated stimuli, such as figure and ground regions, two objects abutting each other, and situations at the border where one object partially occludes another. Previous neurophysiological studies on the segmentation of spatially separated stimuli have mainly focused on the interaction between the RF center and surround (Allman et al., 1985; Born, 2000; Huang et al., 2007, 2008). Britten and Heuer (1999) investigated the spatial summation of multiple stimuli moving in the same direction within the RFs of MT neurons. However, relatively little is known about the neural representation and segregation of multiple stimuli moving differently within the RFs, and more generally, how the visual system uses both motion and spatial cues to segregate multiple stimuli.

Motion discontinuity at the border between two spatially separated stimuli provides key information for segmentation (Nakayama & Loomis, 1974; Regan & Beverley, 1984; Black & Fleet, 2000; Feldman & Weinshall, 2008; Yoonessi & Baker, 2011). In this study, we investigated how MT neurons represent two spatially separated stimuli moving in different directions with a motion border within the RFs. An MT neuron’s spatial RF is approximated by a two-dimensional (2D) Gaussian, with peak sensitivity near the center (Raiguel et al., 1995; Kumano & Uka, 2010). Some MT neurons also exhibit a substructure with multiple distinct hotspots of sensitivity within the RFs (Livingstone et al., 2001; Richert et al., 2013). When two spatially separated stimuli are placed in the RF, according to the neuron’s spatial location selectivity, the neuron may respond to one stimulus more strongly than the other when each stimulus is presented alone, exhibiting a “spatial preference”. It has been shown that neuronal responses in MT to multiple stimuli can be explained by the divisive normalization framework (Carandini & Heeger, 2012), as the combination of the responses elicited by individual stimuli, weighted by their stimulus strengths (Britten & Heuer, 1999; Ni et al., 2012; Xiao et al., 2014; Ni & Maunsell, 2017). However, previous studies have not investigated how the 2D spatial preference of the neurons influences the representation of multiple moving stimuli.

We hypothesized that MT neurons leverage their spatial preferences for individual stimuli within the RFs to better represent and segregate multiple moving stimuli. To test this hypothesis, we placed the motion border between two spatially separated random-dot patches within the RFs. The two stimuli moved in two directions separated by an angle of either 60° or 30°, which was significantly smaller than the mean tuning width (95º-102º) of MT neurons in response to single directions (Albright, 1984; Huang & Lisberger, 2009). At these small angular separations (AS), equal pooling of the responses elicited by individual stimulus components creates a challenging situation for segregating motion directions (Treue et al., 2000; Xiao & Huang, 2015), providing an opportunity to reveal the role of spatial cues in motion segmentation.

We found that the responses of MT neurons to multiple, spatially separated moving stimuli can be well described by weighting the responses to individual stimulus components according to the neurons’ spatial-location preferences for individual stimuli within the RFs. These results can be explained by a spatial preference-weighted normalization model, extending the framework of divisive normalization. We also suggested a circuit implementation of the model. Our results indicate that visual neurons leverage spatial selectivity within their RFs to better represent multiple stimuli, and thereby facilitate visual segmentation.

## Results

We investigated how neurons in area MT encode spatially separated stimuli moving simultaneously in different directions within the neurons’ RFs. Specifically, we investigated the role of spatial cues in representing multiple moving stimuli. We extracellularly recorded electrical activity from isolated neurons in the MT of two male rhesus macaques (R and Z) while they performed a fixation task. Our dataset includes recordings from 117 MT neurons (81 from R and 36 from Z).

Our main visual stimuli consisted of two spatially separated random-dot patches, positioned adjacent to each other along a vertical border and moving in two directions separated by either 60° (Fig. 1A, B) or 30° (Fig. 3). The border was roughly centered on the RFs of the recorded neurons. Nevertheless, it was common for the neuronal responses to the left and right patches presented in isolation to be different, indicating neurons’ spatial preference for sub-regions within the RFs. We varied the vector average (VA) direction of the two stimuli, referred to as the bidirectional stimuli, across 360º to characterize the direction tuning curves. We also measured the direction tuning curves to the left and right patches presented alone.

**Figure 1.**
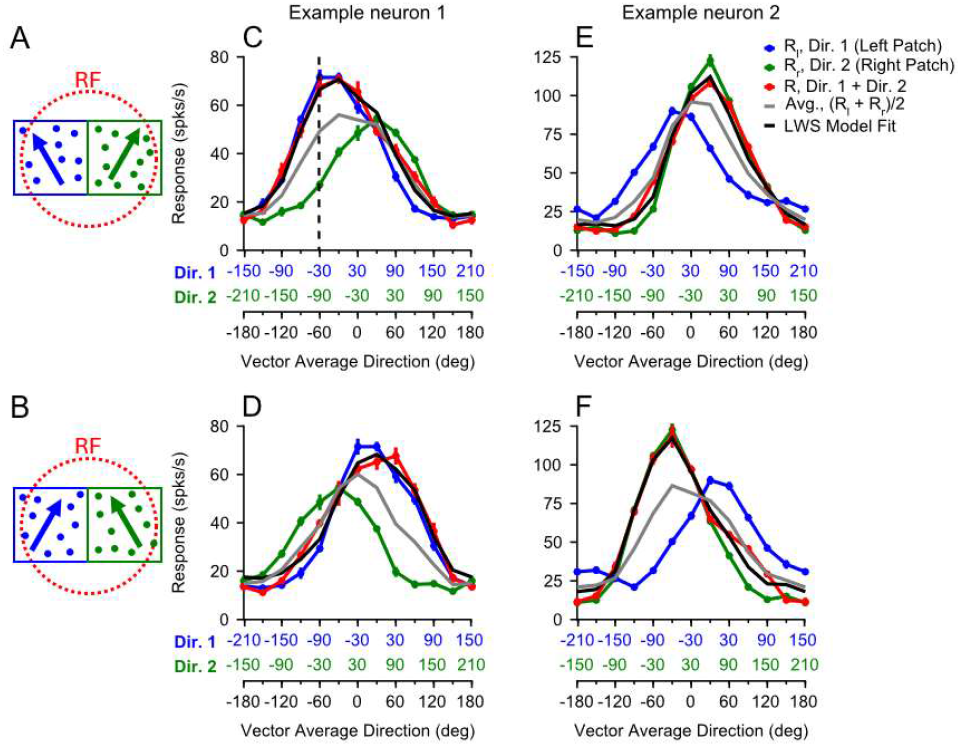
Visual stimuli and response tuning curves of two example neurons. **A, B**. Cartoons depict spatially separated random-dot stimuli placed side by side and two arrangements of motion directions with an AS of 60°. Dots were achromatic. The blue and green colors correspond to the left and right components, respectively. **C**. Response tuning curves from one example neuron. The blue and green abscissae represent the unidirectional components (Dir. 1 and Dir. 2), respectively. The black abscissa corresponds to the VA direction of the bidirectional stimuli. The vertical dashed line demonstrates a VA direction of -60° and the corresponding component directions of -30° (Dir 1.) and -90° (Dir. 2). The black curve is the fit of the LWS model to the bidirectional response (*w*_*l*_= 0.98, *w*_*r*_ = 0.27). **D**. Same neuron as in C with the component directions arranged as shown in B (*w*_*l*_= 0.96, *w*_*r*_ = 0.38). **E, F**. Response tuning curves from another example neuron showing a strong spatial bias toward the right patch, regardless of the direction arrangements (E: *w*_*l*_= 0.26, *w*_*r*_ = 0.98; F: *w*_*l*_= 0.23, *w*_*r*_ = 0.94). Error bars indicate the standard error (SE).

We have previously found that, in response to overlapping random-dot stimuli, some MT neurons have a preference for the motion direction either at the clockwise (C) or counter-clockwise (CC) side of two motion directions, referred to as the “direction side-bias” (Xiao & Huang, 2015). To control for the direction side-bias and examine the effects of spatial cues, our main visual stimuli included conditions in which the motion direction of the right patch was either at the C side (Fig. 1A) or CC side (Fig. 1B) of the left patch’s direction. This design allowed us to separate the effects of spatial cues from direction side-bias.

### Spatial bias in the response tuning to bidirectional stimuli

Figure 1 shows the direction tuning curves of two representative MT neurons in response to two stimulus patches with an AS of 60º. The blue curve shows responses to the left patch alone, the green curve to the right patch alone, and the red curve to both patches moving simultaneously. The three tuning curves are plotted so that the VA direction of the bidirectional stimuli aligns vertically with the corresponding component directions (see the vertical line in Fig. 1C). The responses to the bidirectional stimuli of the first example neuron showed a strong bias toward the directions of the left patch (Fig. 1C, D), substantially different from the average of the responses elicited by the individual component directions (the gray curve). The response bias occurred regardless of whether the left patch moved in the CC side (Fig. 1A, C) or the C side (Fig. 1B, D) of the two component directions, indicating that the bias cannot be explained by a direction side-bias. Note that this neuron’s responses to the individual patches presented alone showed a preference for the left patch – the blue curve had a higher peak than the green curve. The response bias toward the left patch of the bidirectional stimuli is consistent with the neuron’s spatial preference for the individual patches. Another neuron (Fig. 1E, F) had a spatial preference for the right patch when presented alone. In response to the bidirectional stimuli, the response of the neuron was strongly biased toward the right patch, regardless of whether the direction of the right patch was at the C or CC side of the two component directions (Fig. 1E, F).

The responses to the bidirectional stimuli can be well described by a weighted sum of the responses to the individual stimulus components, referred to as the linear weighted summation (LWS) model:

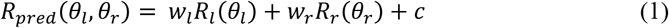

The model has three free parameters. *w*_*l*_ and *w*_*r*_ are the response weights for the left and right components, respectively; *c* is an offset term. *θ*_*l*_ and *θ*_*r*_ are the component directions of the left and right patches, and *R*_*l*_ and *R*_*r*_ are the measured responses to the direction components, respectively. The model-predicted response to the bidirectional stimuli *R*_*pred*_ captured, on average, 90.6% of the response variance across the neurons (std = 9.43%, N = 89). The black curves in Figure 1 illustrate the LWS model fits of the two example neurons.

To isolate the effects of the spatial cues from the direction side-bias, we devised a “spatial location bias index”: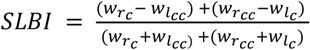, in which *w* is the response weight for a component patch obtained from the LWS model fit of the whole tuning curve of the bidirectional responses. The subscripts *l* and *r* represent left and right patches, and C and CC represent the component direction at the C and CC sides. We compared the weights for the left and right stimuli, *w*_*l*_ and *w*_*r*_, while balancing the direction arrangements. A positive *SLBI* indicates a bias toward the right component patch, whereas a negative *SLBI* indicates a left-patch bias.

Among 89 neurons tested with an AS of 60º, 49 neurons showed a positive *SLBI* (mean ± std: *w*_*r*_ = 0.70 ± 0.15, *w*_*l*_ = 0.34 ± 0.19), and 40 neurons showed a negative *SLBI* (*w*_*r*_ = 0.25 ± 0.20, *w*_*l*_ = 0.77 ± 0.19). Figure 2A-D shows the population-averaged tuning curves of the right- and left-bias neurons. Although the average of the component responses (the gray curve) was only slightly shifted toward the stronger stimulus component, the responses to the bidirectional stimuli (the red curve) showed a strong bias toward the stronger component. The peak of the response tuning curve to the bidirectional stimuli was around the VA directions where one component direction was aligned with the neuron’s preferred direction (PD) (see Fig. 2A), as if the peak response to the bidirectional stimuli was elicited by the PD of one stimulus component without the interference from the other component. Such a representation of the bidirectional stimuli would provide a better segregation of the component directions than averaging the component responses. Of course, the extent of bias in the population-averaged response toward one stimulus component depends on the *SCPI* composition of the neurons in the population.

**Figure 2.**
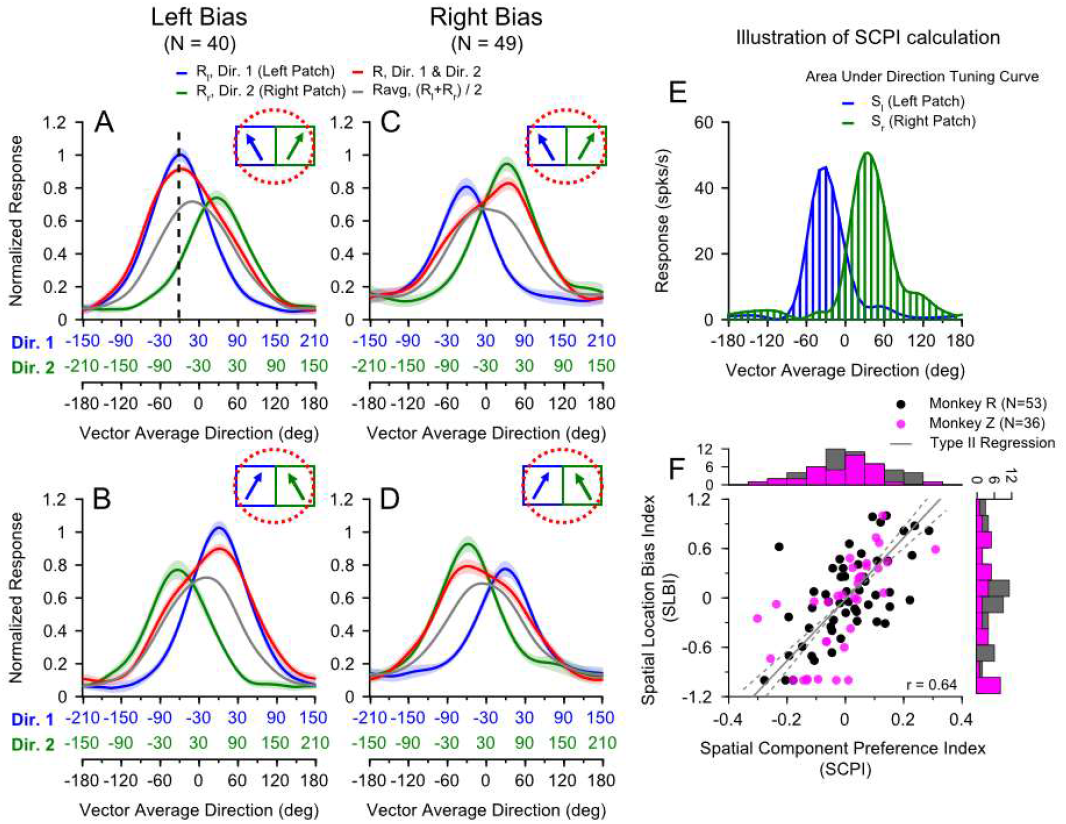
Spatial bias in responses to bidirectional stimuli with an AS of 60°. **A-D**. Spline-fitted and population-averaged response tuning curves. The abscissae, arrangement of stimulus directions, and the colors of the tuning curves follow the same convention as in Figure 1. After spline fitting, the tuning curve of each neuron to the bidirectional stimuli was circularly rotated such that the VA direction of 0° was aligned with the neuron’s PD, before being averaged across neurons. **A, B**. Averaged responses from neurons that show a spatial bias towards the left side of the RF (i.e., *SLBI* < 0) when the left patch moved at the CC side (A), or C side (B) of the two motion directions. The vertical dashed line in A shows a VA direction of -30° and the corresponding component directions of 0° (Dir 1), which is the aligned preferred direction of the neurons, and -60° (Dir. 2). **C, D**. Averaged responses of neurons showing a spatial bias towards the right side of the RF (i.e., *SLBI* > 0). Error bands indicate the SE. In A-D, the tuning curve to the bidirectional stimuli peaked around ±30º VA directions, where one of the component directions was aligned with the neuron’s PD (0º). **E**. Illustration of how *SCPI* was calculated. The blue and green shades depict the areas under the tuning curves (AUC) for the left (*S*_*l*_) and right (*S*_*r*_) stimulus components from an example neuron. **F**. Scatter plot showing the relationship between *SLBI* and *SCPI* across neurons and the marginal distributions. Each dot represents data from one neuron. The solid gray line indicates the best fit from Type II regression, while the dotted lines indicate the lower and upper bounds of the 95% confidence interval (CI) of the regression.

**Figure 3.**
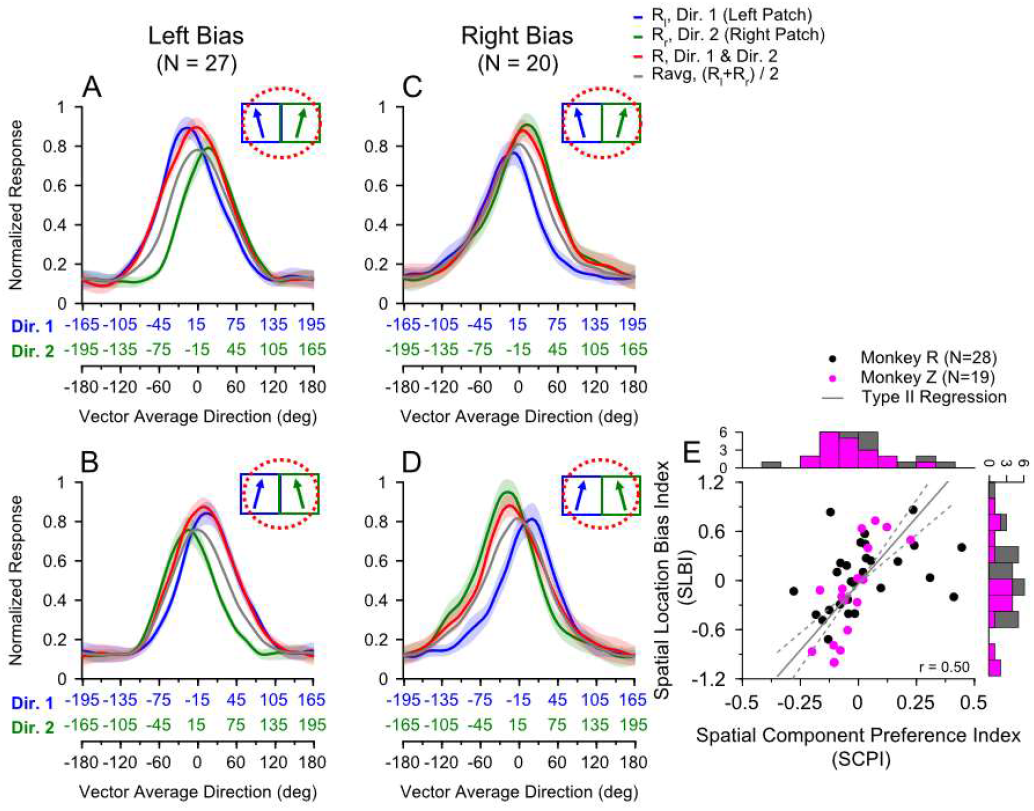
Spatial bias in responses to bidirectional stimuli with an AS of 30°. **A-D**. Spline-fitted and population-averaged response tuning curves. **A, B**. Tuning curves from a population of neurons showing spatial bias towards the left stimulus component (*SLBI* < 0). **C, D**. Neurons showing spatial bias towards the right stimulus component (*SLBI* > 0). **A, C**. The left patch moved in the CC-side direction. **B, D**. The left patch moved in the C-side direction. Error bands indicate the SE. **E**. Scatter plot showing the relationship between *SLBI* and *SCPI* across neurons and the marginal distributions. The solid gray line shows the Type II regression, while the dotted lines indicate the lower and upper bounds of the 95% CI.

To characterize the neuron’s spatial preference for individual stimuli presented alone, we used a “spatial component preference index”: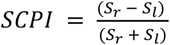, in which *S*_*r*_ and *S*_*l*_ are the integrals of the direction tuning curves to the right and left patches, respectively (illustrated in Fig. 2E). The integral of a tuning curve for a stimulus component corresponds to the summed responses of a neuronal population with identical tuning curve shapes but different preferred directions.

We found that the sign and magnitude of the spatial bias in the neuronal response to the bidirectional stimuli (*SLBI*) were positively correlated with the neurons’ spatial preference for single patches (*SCPI*) (Fig. 2F, Type II regression, *r* = 0.64, 95% CI of the slope: 3.1 to 4.4). While the *SCPI* values were distributed mainly within a narrow range from −0.18 to 0.14 (central 80% relative to the median), the *SLBI* values had a significantly wider spread from -0.99 to 0.66 (central 80%) (Levene’s test for equality of variances, p = 2.05×10^−15^). These distributions suggest that a slight difference in the spatial preference for the stimulus components can lead to a substantial spatial bias in the responses to the bidirectional stimuli.

As the difference between the directions of motion components decreases, segregation of multiple moving stimuli becomes more difficult. We asked whether spatial bias also occurred at smaller AS to facilitate segregation. We recorded from 47 MT neurons using an AS of 30º. Twenty-seven neurons had a negative *SLBI*, and the population-averaged response to the bidirectional stimuli showed a strong bias toward the left patch (Fig. 3A, B). Twenty neurons had a positive *SLBI*, and the population-averaged response to the bidirectional stimuli showed a strong bias toward the right patch (Fig. 3C, D). The LWS model also provided a good fit to the response tuning curves, with a mean variance accounted for of 95.1% (std = 5.5%, N = 47). Similar to the AS of 60º, the *SLBI* for the AS of 30º was positively correlated with the *SCPI* (Type II regression, *r* = 0.50, 95% CI of the slope: 2.4 to 4.1), and the variances of *SLBI* and *SCPI* are significantly different (Levene’s test, p = 9.96×10^−10^) (Fig. 3E). The spatial bias allowed neurons to represent the motion component at their preferred spatial location more effectively than would be possible if they simply averaged (or equally pooled) the responses to the stimulus components. For a population of neurons with different spatial preferences, different component directions with a small AS can be represented by different neurons, thereby facilitating segregation.

### Properties of the spatial bias

In the following two experiments, we used bidirectional stimuli with an AS of 60º to further characterize the properties of the spatial bias. First, we asked whether the responses of V1 neurons with RFs at the motion border contributed to the spatial bias in MT. With a sharp border between two moving stimuli as used in our main experiments, V1 neurons with RFs at the border can be stimulated by both stimulus components. A previous study from our lab suggested that normalization in V1 can have a strong effect on MT responses to multiple stimuli (Wiesner et al., 2020). To address this question, we placed a 1.5° gap between the two stimulus patches (Fig. 4C). At the eccentricities we recorded from MT neurons (mean = 9.3°, std = 4.4°), V1 RFs are typically smaller than 1.5° (Dow et al., 1981). We recorded from 20 neurons under both gap (Fig. 4C, D) and no-gap (Fig. 4A, B) conditions. The response tuning and the spatial bias were very similar between the two conditions. Twelve neurons showed a left bias (Fig. 4A, C), and eight neurons showed a right bias (Fig. 4B, D). The *SLBI* values from gap and no-gap conditions were not significantly different (p = 0.42, Wilcoxon Signed Rank test) (Fig. 4E). These results suggest that the spatial bias did not originate from V1 neurons with RFs at the motion border.

**Figure 4.**
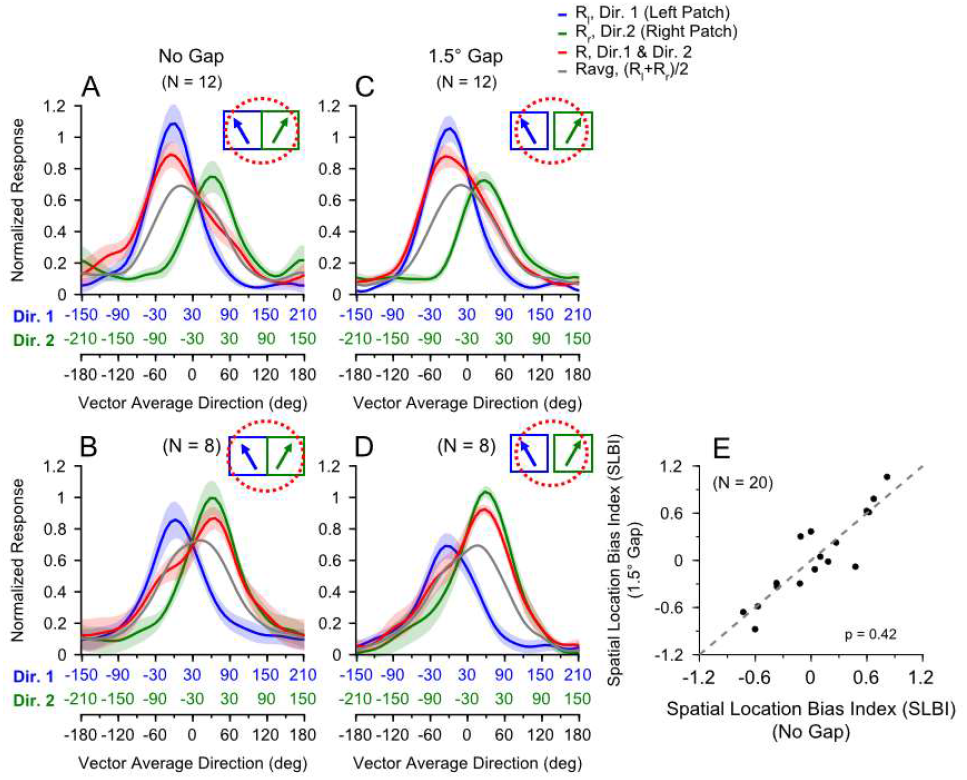
Consistent spatial biases with and without a gap between the two stimulus patches. The same group of neurons (N = 20) was tested with both stimulus conditions. The left patch always moved in the CC-side direction, and the right patch moved in the C-side direction. **A, B**. Population-averaged direction tuning curves without the gap. **C, D**. With a 1.5° gap. The green and blue abscissae follow the same convention as in Figure 1. Error bands indicate the SE. **E**. Scatter plot of the *SLBI* with and without the gap. Each dot represents *SLBIs* from one neuron. The gray dashed line is the unity line. We calculated 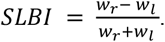 *w*_*r*_ and *w*_*l*_ are the weights for the right and left patch, respectively, fitted by the LWS model.

We have shown that the spatial bias in the response to bidirectional stimuli (measured by *SLBI*) is correlated with the spatial preference for the individual stimulus components (measured by *SCPI*). To further understand the relationship between *SCPI* and *SLBI*, in the second experiment, we manipulated the spatial preference of the neurons for the stimulus components by horizontally shifting the position of the motion border between the two stimulus patches. We recorded from 13 neurons, testing each with at least two positions. Figure 5A, B shows the direction tuning curves of an example neuron in response to stimuli with a motion border placed at two locations. At one border position, the neuron showed a spatial preference for the left patch (*SCPI* = -0.19) and its response tuning to the bidirectional stimuli showed a left bias (*SLBI* = - 0.23) (Fig. 5A). When we shifted the motion border, the same neuron showed a spatial preference for the right patch (*SCPI* = 0.25) and response tuning to the bidirectional stimuli showed a strong right bias (*SLBI* = 0.63) (Fig. 5B). The change of *SLBI* as we varied the *SCPI* was highly consistent across the neurons (Fig. 5C). The change of the *SLBI* between two positions was highly correlated with the change of the *SCPI* of the same neuron (Pearson correlation coefficient *r* = 0.95, p < 0.0001) (Fig. 5D). These results suggest that the spatial bias in the responses to multiple spatially separated stimuli is dependent on the spatial preference of the neuron for the stimulus components.

**Figure 5.**
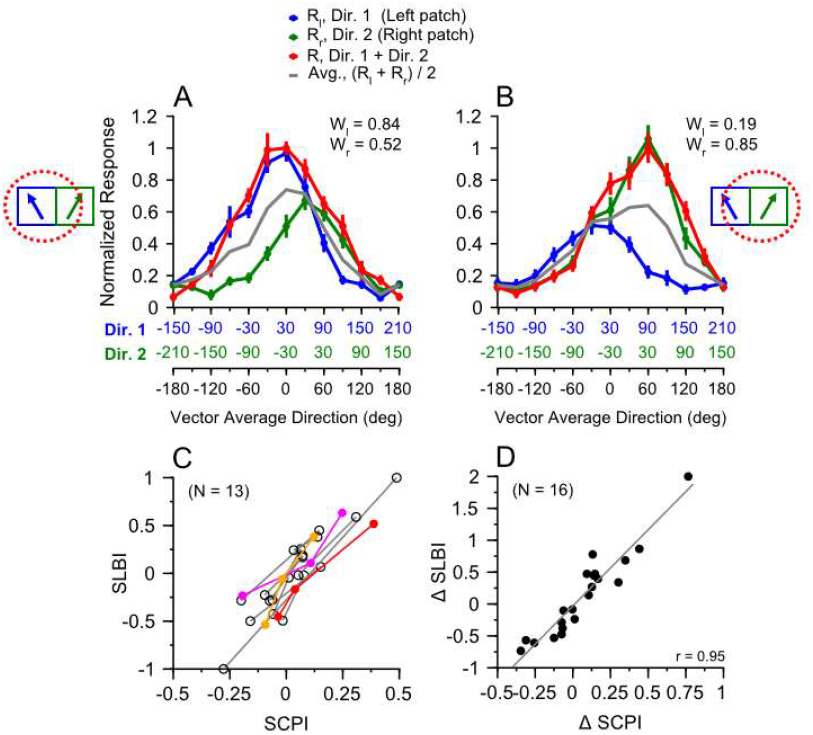
Changing a neuron’s spatial preference for the stimulus components by shifting the border position led to a corresponding change in the spatial bias. In this experiment, the left patch always moved in the CC-side direction, and the right patch moved in the C-side direction. 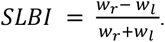 **A, B**. Response tuning curves of one example neuron to visual stimuli and motion border placed at different locations. In A, the left patch was closer to the center of the RF, whereas in B, the right patch was closer to the center. The weights *w*_*r*_ and *w*_*l*_ were fitted by the LWS model. Error bars represent the SE. **C**. Results from 13 neurons showing how the *SLBI* changed with the *SCPI*. Each line connects two motion border positions from the same neuron. Three neurons were tested with three border positions (illustrated by red, magenta, and orange colors). The remaining neurons were tested with two border positions (black). **D**. The change of the *SLBI* between two border positions was highly correlated with the change of the *SCPI*. Each dot represents the Δ*SLBI* and the Δ*SCPI* between two positions from one neuron. Each of the three neurons tested with three positions contributed two dots (between positions 1 and 2 and between 1 and 3). The black line shows the linear regression fit.

### Temporal development of the spatial bias

To provide insight into the neural mechanism underlying the spatial bias, we characterized the time course of the neuronal responses to bidirectional stimuli with an AS of 60°, using the data set shown in Figure 2. We measured the direction tuning curves with a 50-ms time window, sliding in a 10-ms step. We pooled the tuning curves (with flipping as needed) to bidirectional stimuli that showed left and right biases (as illustrated in Fig. 2A-D), such that the spatial biases were all aligned with the negative VA directions and the potential effect of direction side-bias was averaged out (see Methods). The initial response tuning to the bidirectional stimuli (*R*) showed no spatial bias at the time window centered on 45 ms (Fig. 6A, B). The spatial bias began at ∼55 ms and developed over the next 90 ms before reaching its peak (Fig. 6A, B, E). The response peak of the tuning curve to the bidirectional stimuli eventually reached the VA direction of about -30°, where one of the component directions was at the neuron’s PD (i.e., 0°) (Fig. 6E, red curve). In other words, over time, the response peak to the bidirectional stimuli was aligned with the response peak to the component direction at the biased side, thereby facilitating the representation of the component direction. In contrast, the average of the direction tuning curves to the individual stimulus components (*R*_*avg*_) had a slight shift toward the stronger component response (Fig. 6C, D, E). This shift was relatively stable over time, and the response peak was around the VA direction of about -10° (Fig. 6E, black curve), much smaller than the actual response bias to the bidirectional stimuli.

**Figure 6.**
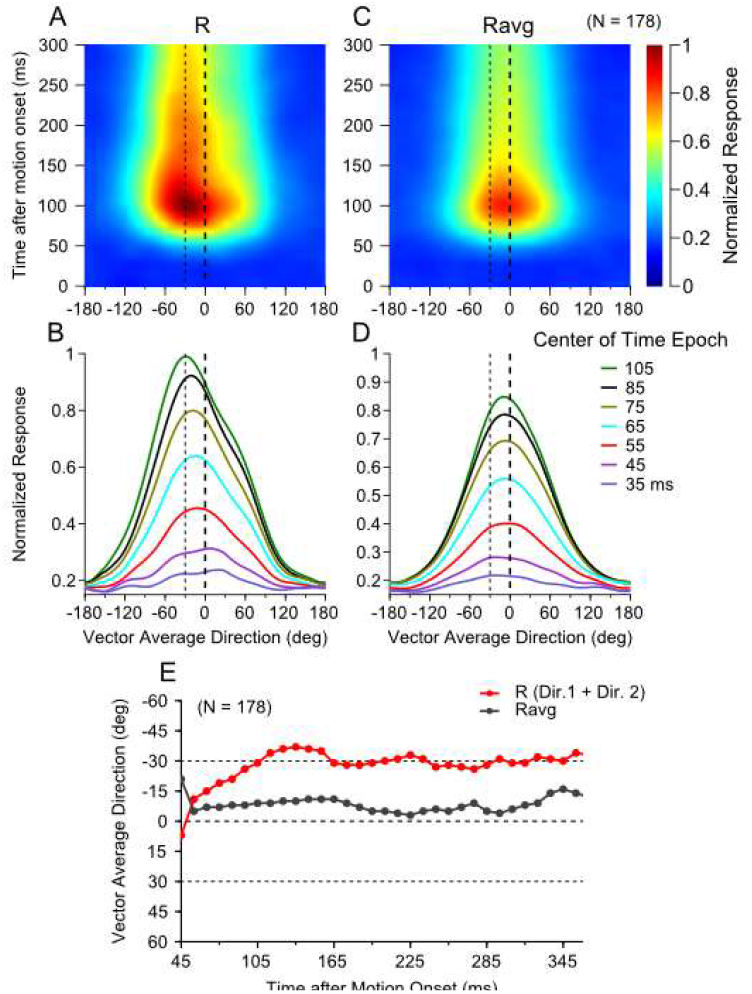
Time course of the neuronal response to the bidirectional stimuli. Each neuron contributed to two tuning curves with one patch moving in either the CC- or C-side of two directions (a total of 178 tuning curves from 89 neurons). The tuning curves were aligned (and flipped relative to VA 0° as needed), such that the spatial bias of each neuron was always toward the negative VA directions. **A, B**. Time course of the population-averaged response to the bidirectional stimuli. The thick dashed line marks VA 0°. The thin dotted line shows the VA direction of -30°, at which one of the stimulus components moved in the PD of the neuron. The ordinate in A indicates the center of each time epoch, relative to the motion onset. **C, D**. Time course of the average of the responses elicited by individual stimulus components. **E**. Time course of the response peak of the tuning curves to the bidirectional stimuli (red) and the averaged component responses (black). The ordinate shows the peak location relative to the VA direction 0°. The abscissa shows the center of each time epoch after motion onset. The dashed line at -30° indicates the VA direction at which one stimulus component was moving in the PD of the neuron.

### Responses to spatially separated stimuli captured by spatial preference-weighted normalization

Our results demonstrate that neuronal responses to spatially separated bidirectional stimuli can be described by a weighted sum of the neuron’s responses to the individual stimulus components. A central question, then, is what determines these weights. The divisive normalization framework (Carandini & Heeger, 2012) has provided a powerful account of diverse neural phenomena, including responses to multiple visual stimuli. In this model, the role of the denominator, which scales activity by the population response of the normalization pool, is well established. However, the nature of the numerator is less understood. We have previously proposed that the weight associated with a given stimulus component is proportional to the activity of a “weighting pool” elicited by the stimulus component (Xiao et al., 2014; Wiesner et al., 2020; X. Huang et al., 2025). The composition of the weighting pool remains unclear.

We extended the normalization model by assuming that the weighting for a stimulus component reflected a neuron’s spatial preference for that component. We approximated the spatial preference of an MT neuron by summing the neuron’s direction tuning curve across 360º to each stimulus component. The summed response was invariant to the direction of the stimulus component and used as the numerator in the extended normalization equation (Eq. 2):

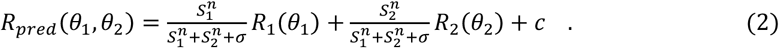

*R*_*pred*_ is the model-predicted response to the bidirectional stimuli. *R*_1_ and *R*_2_ are the recorded responses to the individual stimulus components. *θ*_1_ and *θ*_2_ are the two component directions. *S*_1_ and *S*_2_ are the summed responses of the direction tuning curves to the individual stimulus components:

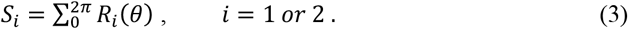

The model has three free parameters *n, σ*, and *c*, with the constraints of 0.01 ≤ *n* ≤ 100, and 0 ≤ *c* ≤ 100. For a neuron that prefers stimulus component 1 over 2, *S*_1_ is greater than *S*_2_. Therefore, the weight for *R*_1_ is greater than that for *R*_2_, making the spatial bias consistent with the neuron’s spatial preference. The power-law function and divisive normalization in Eq. 2 contribute to the nonlinearity of the response weight.

This extended normalization model provided a good fit of the whole tuning curve to the bidirectional stimuli, accounting for on average 86.3% (std = 12.4%, N = 89) of the response variance for the AS of 60°, and 93.7% (std = 5.9%, N = 47) of the response variance for the AS of 30°. In addition, the exponent parameter (*n*) in the normalization model fit had a median value greater than 1 (4.4 for AS 60°, and 7.5 for AS 30°), indicating an expansive power law, which explains why a slight change in spatial preference could lead to a substantial change in spatial bias.

The goodness-of-fit of the normalization model is comparable to that of the LWS model (90.6 ± 9.4% for AS 60°, and 95.1 ± 5.5% for AS 30°, Fig. 7A, B), although the LWS model fits were significantly better (t-test, p = 8.2×10^−7^ for AS 60°, p = 5.16×10^−5^ for AS 30°). This difference is expected because the weights in the LWS are free parameters, whereas the weights in the normalization model must follow a prescribed formula based on the neuron’s spatial preference. The weights for the stimulus components, and therefore the *SLBI*, derived from the extended normalization model were in close agreement with those derived from the LWS model (Fig. 7C, D, Wilcoxon signed-rank test, p = 0.75 and p = 0.40, respectively). These results provide a computational explanation for the spatial bias observed in responses to bidirectional stimuli.

**Figure 7.**
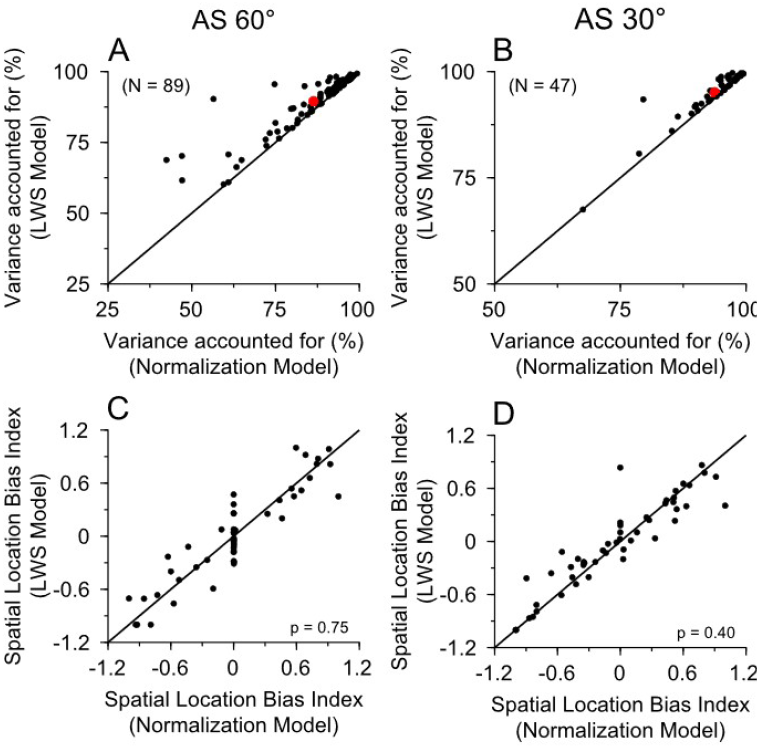
Comparison of the goodness of fit and *SLBI* between the normalization and LWS model. Each dot represents a single neuron. **A, B**. Scatter plot comparing the variance accounted for by the LWS (ordinate) and the normalization (abscissa) model for AS of 60° (A) and 30° (B). The red dot indicates the population means. **C, D**. Comparison of *SLBI* calculated using the weights from the LWS and normalization model fits, for AS of 60° (C) and 30° (D).

We proposed a circuit implication for the spatial preference-weighted normalization, as illustrated in a schematic diagram (Fig. 8). We assumed that two populations of direction-selective neurons (P and Q) in V1 have RFs covering the two spatially separated stimulus components, respectively. Neurons in P and Q project to an MT neuron *j*, as well as other MT neurons that have the same RF location and the same spatial preference as MT neuron *j*, but with different direction preferences. At each spatial location (circle within the shaded blue and green ovals), neurons in P and Q also have PDs spanning 360°. *C*_*ij*_ is the synaptic weight between neuron *i* in population P (or Q) and the MT neuron *j. C*_*ij*_ is subject to gain control.

**Figure 8.**
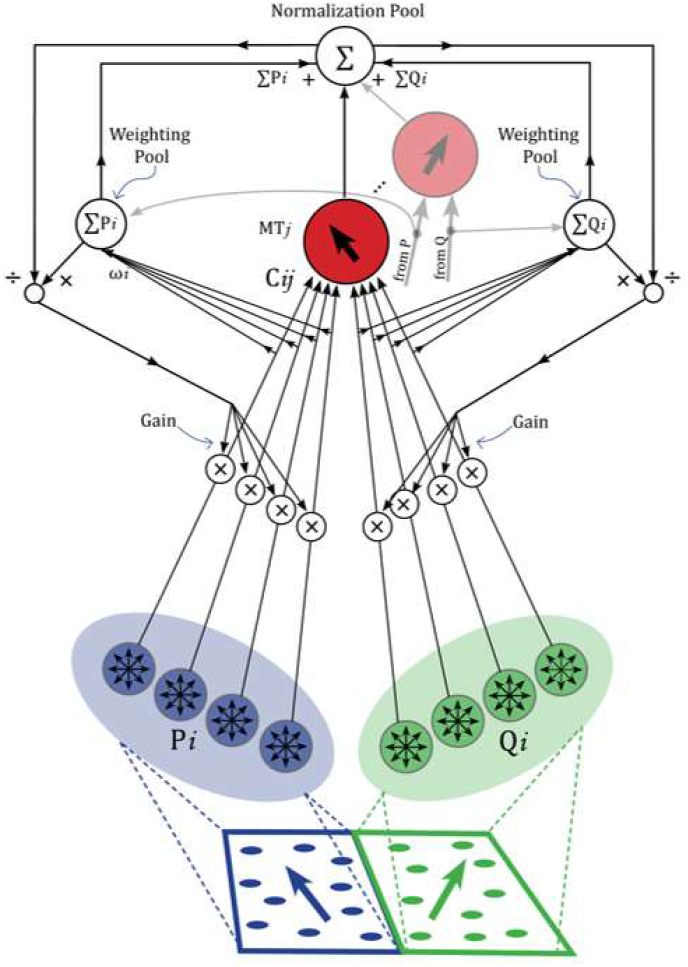
A schematic circuit diagram of the spatial preference-weighted normalization model. An MT neuron *j* (red circle) receives inputs from two populations of direction-selective neurons (P and Q), presumably in V1. Each population (P or Q) contains neurons with a broad range of direction preferences (black arrows). Neurons in population P have RFs covering one stimulus component (blue), and Q covering the other stimulus component (green). *C*_*ij*_ is the synaptic weight between neuron *i* in P (or Q) and the MT neuron *j*. The activity of the weighting pool includes the summed response from the inputs to MT neuron *j*, as well as to other MT neurons that have the same RF location and the same spatial preference as MT neuron *j*, but with different direction preferences across 360°. The activity of the normalization pool is the sum of the activities of the two weighting pools, as well as the responses of a population of MT neurons. The gain factor for *C*_*ij*_ is controlled by multiplying the response of the weighting pool for P or Q, respectively, and dividing by the activity of the normalization pool.

The activity of the weighting pool includes the sum of the inputs from neurons in P (or Q) projecting to a group of MT neurons that have different direction preferences across 360°, but have the same RF location and the same spatial preference as MT neuron *j*. As a result, the summed activity in the weighting pool reflects an MT neuron’s spatial preference for a stimulus component, regardless of the stimulus direction or the direction preference of the MT neuron.

The activity of the normalization pool is the sum of the activities of the two weighting pools, as well as the responses of a population of MT neurons. The gain factor *g* for *C*_*ij*_ is controlled by multiplying the response of the weighting pool for inputs from P or Q, respectively, and dividing by the response of the normalization pool.

## Discussion

We found that MT neurons showed a spatial-location bias in their responses to multiple, spatially separated stimuli moving within their RFs, and this bias was dependent on the neuron’s spatial preference for the individual stimuli. The sign and magnitude of the spatial bias in response to multiple stimuli were positively correlated with the neurons’ spatial preference. Additionally, a small difference in responses to individual stimuli at different locations could lead to a substantial spatial bias. We also found that the spatial bias developed over time, reaching its maximum approximately 90 ms after its onset. We demonstrated that MT neuronal responses to multiple stimuli can be well described by a sum of the responses elicited by individual stimuli, weighted by the neuron’s spatial preference for the stimulus components, and the response weights are invariant to the stimulus directions.

### The spatial bias cannot be explained by attention modulation

Spatial attention directed to one of multiple stimuli within MT neurons’ RFs can significantly modulate the neuronal response (Treue & Maunsell, 1996; Ferrera & Lisberger, 1997; Treue & Maunsell, 1999; Seidemann & Newsome, 1999; Lee & Maunsell, 2010). However, the spatial bias found in this study was unlikely to be contributed by spatial attention for the following reasons. First, some of our recordings were conducted using a multi-channel electrode array. We found that simultaneously recorded neurons could exhibit left- and right-biases in the same sessions, which cannot be explained by an attention modulation. Second, we collected data from 20 neurons from monkey R in response to our main stimuli within the RFs with an AS of 60°, while the monkey performed a direction discrimination task with attention directed away from the RFs to the visual hemifield opposite to the RFs (see Methods). The behavior task engaged the monkey’s attention and was demanding. The monkey performed the task well, with a mean correct rate of 77.4% (std = 7.9%) in 18 recording sessions. Using this attention-away task, we found that the *SLBI* were positively correlated with the *SCPI* (Type II regression, *r* = 0.56), consistent with the results obtained with the fixation task.

### Spatial preference-weighted representation of multiple stimuli

The well-accepted divisive normalization framework (Carandini & Heeger, 2012) has suggested that the neuronal response to multiple stimuli is weighted by the signal strengths of the individual stimuli, such as the luminance contrast (Busse et al., 2009; MacEvoy et al., 2009; Reynolds & Desimone, 2003; Bao & Tsao, 2018; Heuer & Britten, 2002; Ni et al., 2012; Xiao et al., 2014; Ni & Maunsell, 2017) and motion coherence (Morgan et al., 2008; Xiao et al., 2014). However, previous studies have not demonstrated the role of the spatial selectivity of neurons in representing multiple stimuli. Our study helps to establish that the neuronal response to multiple stimuli is weighted by neurons’ 2D spatial preferences for individual stimuli within the RFs. This finding made an important extension of the divisive normalization model and revealed a key mechanism for representing and segregating spatially separated stimuli.

A recent report has shown a similar result that MT responses to two moving stimuli follow a normalization model weighted by the product of the stimulus contrast and the RF weight for spatial location (Cherian & Maunsell, 2025). Our lab has also demonstrated that neuronal responses in MT to two overlapping stimuli moving at different depths (binocular disparities) exhibit a bias toward one of the binocular disparities, and this disparity bias is positively correlated with the neuron’s disparity preference for single stimuli (Chakrala et al., 2024). Since disparity selectivity can be considered a form of spatial selectivity in depth, spatial preference-weighted representation of multiple stimuli may be general to neurons’ selectivity for locations in three-dimensional space.

### Neural mechanisms underlying spatial preference-weighted normalization

We previously suggested that MT responses to two moving stimuli with competing signal strengths can be governed by response normalization in V1 when individual V1 neurons are driven by both (superimposed) stimuli, but not when V1 neurons are driven by only one stimulus component of spatially separated stimuli (Wiesner et al., 2020). Our finding of similar spatial bias in MT using the stimuli with a gap (Fig. 4) suggests that the spatial bias in MT was not simply inherited from response normalization in V1.

Although our model (Eq. 2) is descriptive, it provides several insights into possible circuit mechanisms of spatial preference-weighted normalization (Fig. 8). First, the model suggests that weighting by spatial preference is applied to the response elicited by each stimulus component. One way to implement this is to pool the inputs to the MT neuron that represent the stimulus component, and the weighting also occurs at the inputs to the MT neuron in a form of gain control. Second, our model and data suggest that the weights should be invariant to stimulus directions. Therefore, we propose that the weighting pool should include activities from neurons with different PDs spanning 360°. This can be implemented by equally pooling the inputs from V1 neurons that have different PDs to the MT neuron. Third, the weighting should reflect the spatial preference, rather than the direction preference of the MT neuron. The synaptic connections from a population of V1 neurons to an MT neuron may preserve both spatial selectivity and direction selectivity. One way to implement this is to pool across the synaptic inputs from the same V1 population (e.g., Q in Fig. 8) projecting to different MT neurons that have the same RF location and the same spatial preference, but with different direction preferences. By pooling across the inputs to these MT neurons, the pooled activity preserves the spatial preference of the MT neuron, but is invariant to the direction selectivity of the MT neuron. Finally, our model distinguishes the weighting pool from the normalization pool, which have different compositions (Fig. 8).

Our timecourse analysis showed that the spatial bias developed over ∼90 ms, suggesting that spatial bias may arise not only from feedforward pooling but also from recurrent interactions between neurons. Clarifying these mechanisms is a worthwhile direction for future work.

### Spatial bias and strategy for segregating multiple stimuli

Our results provide a critical insight into how the visual system represents and segregates multiple visual stimuli. By leveraging the spatial selectivity within the RFs, the visual system nonlinearly amplifies selectivity for single stimuli at different locations, resulting in a substantial spatial bias in the response to multiple stimuli. In this way, neuronal responses to multiple stimuli within the RFs are not equally mixed, but mainly represent the stimulus located at the more preferred location of the neurons. For a population of neurons with various spatial preferences, this coding strategy would enable individual components of multiple stimuli to be represented by different neurons within the population, thereby facilitating the segregation of multiple stimuli. This spatial location-based segregation is particularly useful when other features of the visual stimuli differ only slightly, such as motion directions with a small angular separation in our stimuli. Theoretical studies suggest that, rather than mixing multiple stimuli with equal weight, random mixing with different weights by individual neurons can significantly improve the ability to encode multiple stimuli in neuronal populations (Orhan & Ma, 2015; Huang et al., 2017). The spatial bias found in this study demonstrates that the visual system takes this theoretically effective strategy to represent and segregate multiple visual stimuli.

## Materials and Methods

### Subjects and neural recording

Two male adult rhesus monkeys (*Macaca mulatta*) participated in the experiments. Experimental protocols were approved by the Institutional Animal Care and Use Committee of UW-Madison and conform to U.S. Department of Agriculture regulations and to the National Institutes of Health guidelines for the care and use of laboratory animals. A headpost and a recording cylinder were implanted during sterile surgery with the animal under isoflurane anesthesia. We took a vertical approach to access area MT. We identified MT by its characteristically large portion of directionally selective neurons, small RFs relative to those in the neighboring area MST, its location at the posterior bank of the superior temporal sulcus, and visual topography of the RFs (Gattass & Gross, 1981). For the majority of the electrophysiological recordings, we used tungsten electrodes (1-3 MΩ, FHC). For some experiments, we used a 16-channel linear electrode array (S-Probe, Plexon) to simultaneously record multiple neurons. Electrical signals were amplified, and single units were identified with a real-time template-matching system and an offline spike sorter (Plexon). Eye position was monitored using a video-based eye tracker (EyeLink, SR Research) with a rate of 1000 Hz.

### Visual stimuli and experimental procedure

Stimulus presentation and data acquisition were controlled by a real-time data acquisition program “Maestro” (https://sites.google.com/a/srscicomp.com/maestro/home). Visual stimuli were presented on a 25-inch CRT monitor at a viewing distance of 63 cm. Monitor resolution was 1024 × 768 pixels (or 1152 × 864 pixels) with a refresh rate of 100 Hz. Stimuli were generated by a Linux workstation using an OpenGL application. Visual stimuli were achromatic random dots moving within static square apertures. In the initial data collection, each random-dot patch was 10° × 10°. For most of the data collection, each patch was 5° × 5°. Each dot was a tiny square extending ∼0.08°. The dots had a luminance of 22 cd/m^2^ presented on a uniform background of 10 cd/m^2^, resulting in a Michelson contrast of 0.375. The dot density was 2.7 dots/deg^2^. Random dots in each patch moved in a specified direction, with a motion coherence of 100%. The lifetime of each dot was as long as the motion duration. If a dot reached the boundary of the aperture during the motion period, it was immediately replotted on the other side of the aperture and kept moving.

In each experimental trial, the monkey maintained fixation within a 1° × 1° electronic window around a small fixation point. After an MT neuron was isolated, we characterized its direction selectivity, preferred speed, and RF location using methods described previously (Wiesner et al., 2020). In our main experiment, the visual stimuli were presented after the monkey maintained fixation for 250 ms. To isolate the response to motion from that elicited by the stimulus onset, the visual stimuli were first turned on and remained stationary for 300 ms before moving for 500 ms. The visual stimuli were then turned off. The monkey maintained fixation for an additional 250 ms. To characterize the response tuning curve to bidirectional stimuli, we varied the VA direction around 360° in steps of either 30° (for an AS of 60°) or 15° (for an AS of 30°). We also characterized the direction-tuning curves for individual stimulus patches presented alone at different spatial locations. The trials presenting bidirectional stimuli and individual stimulus patches were randomly interleaved.

To control for spatial attention, in a subset of experiments, we recorded from MT neurons while one monkey (R) attended a stimulus in the visual hemifield opposite from the RFs of the recorded MT neurons. The attended stimulus consisted of random dots moving within a stationary circular aperture with a diameter of 5°, placed 10° to the left of the fixation point. The random dots moved in a single direction, on either the C or CC side of the upward direction, offset by 10°, 15°, or 20°. The monkey performed a direction discrimination task, reporting whether the stimulus moved in a direction at the C or CC side of the upward direction. While the monkey performed this task, our main visual stimuli (the bidirectional stimuli and the stimulus components) were placed in the RFs as in the fixation task. After 200 ms of fixation, both the attended stimulus and the RF stimulus were turned on simultaneously and remained stationary for 250 ms before moving for 400 ms. After the motion period, both stimuli were turned off, and two reporting targets located on the left and right sides of the fixation point were turned on. The animal made a saccadic eye movement to one of the two reporting targets to indicate the choice. Correct trials were rewarded with juice. All trials were randomly interleaved.

### Data analysis

Response firing rate was calculated during the period of stimulus motion and averaged across repeated trials. The raw direction tuning curves for the bidirectional stimuli and the individual stimulus components were spline-fitted using the Matlab function *csaps* with a smoothing parameter *p* set to 0.009 and a step of 1°. We then rotated the spline-fitted tuning curve to the bidirectional stimuli such that the VA direction of 0° was aligned with the PD of each neuron. The responses of each neuron to the bidirectional stimulus and individual stimulus components were normalized by the maximum spline-fitted response to the bidirectional stimuli. We then averaged the aligned and normalized tuning curves across neurons to obtain population-averaged tuning curves. To determine a neuron’s spatial preference for the stimulus components, we calculated the areas under the response tuning curves (AUC) for the individual stimulus components by summing the firing rates of the spline-fitted tuning curves across 360°.

To characterize the timecourse of the spatial bias, we combined direction tuning curves showing either left or right bias, and toward either the C and CC sides of the component directions. To achieve this, we flipped the response tuning curves when the spatial bias was toward the C side directions, relative to the VA direction of 0°. This flip aligned the response bias toward the CC side directions (i.e., negative VA directions). For the tuning curves that were biased toward the CC side directions, we did not flip them. We then averaged all the flipped and not-flipped tuning curves across the neurons, with the spatial bias always toward the negative VA directions.

Type II regression fitting was done using the *gmregress* function in Matlab created by Trujillo-Ortiz and Hernandez-Walls (2010). The function is based on the model description and implementation from Ricker (1973).

### Model fit of direction tuning curves

To fit the direction tuning curves of the spatially separated stimuli moving in different directions, we used a linear weighted summation model (Eq. 1) as well as an extended divisive normalization model (Eq. 2). These model fits were derived using the constrained minimization tool “fmincon” (MATLAB) to minimize the sum of squared error. To evaluate the goodness of fit of a model for the response tuning to the bidirectional stimuli, we calculated the percentage of variance (PV) accounted for by the model as:

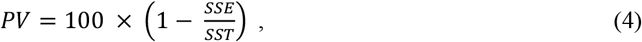

where SSE is the sum of squared errors between the model fit and the neuronal data, and SST is the sum of squared differences between the data and the mean of the data.

## Acknowledgments

We thank Bryce Arseneau for animal training, Drs. Jennifer Coonen and Kevin Brunner at the Wisconsin National Primate Research Center for excellent veterinary care and surgical assistance.

## Funding support

This research was supported by the National Eye Institute Grant R01 EY022443. B.G. was partially supported by a Walsh Graduate Student Support Initiative award and Kenzi Valentyn Vision Research award from the McPherson Eye Research Institute at the University of Wisconsin–Madison.

## Notes

### Competing Interest Statement

The authors have declared no competing interest.

### Summary of Updates

This version added additional analysis results (Figure 5) and revised the text of the manuscript.

